# Biochemical Validation of a Fourth Guanidine Riboswitch Class in Bacteria

**DOI:** 10.1101/2020.10.21.349134

**Authors:** Hubert Salvail, Aparaajita Balaji, Diane Yu, Adam Roth, Ronald R. Breaker

**Affiliations:** Department of Molecular, Cellular and Developmental Biology, Yale University, New Haven, Connecticut 06520-8103, USA; Howard Hughes Medical Institute, Yale University, New Haven, CT 06520-8103, USA; Department of Molecular Biophysics and Biochemistry, Yale University, New Haven, Connecticut 06520-8103, USA

## Abstract

An intriguing consequence of ongoing riboswitch discovery efforts is the occasional identification of metabolic or toxicity response pathways for unusual ligands. Recently, we reported the experimental validation of three distinct bacterial riboswitch classes that regulate gene expression in response to the selective binding of a guanidinium ion. These riboswitch classes, called guanidine-I, -II and -III, regulate numerous genes whose protein products include previously misannotated guanidine exporters and enzymes that degrade guanidine via an initial carboxylation reaction. Guanidine is now recognized as the primal substrate of many multidrug efflux pumps that are important for bacterial resistance to certain antibiotics. Guanidine carboxylase enzymes had long been annotated as urea carboxylase enzymes but are now understood to participate in guanidine degradation. Herein we report the existence of a fourth riboswitch class for this ligand, called “guanidine-IV”. Members of this class use a novel aptamer to selectively bind guanidine and use an unusual expression platform arrangement that is predicted to activate gene expression when ligand is present. The wide distribution of this abundant riboswitch class, coupled with the striking diversity of other guanidine-sensing RNAs, demonstrates that many bacterial species maintain sophisticated sensory and genetic mechanisms to avoid guanidine toxicity. This finding further highlights the mystery regarding the natural source of this nitrogen-rich chemical moiety.

---

Numerous riboswitch classes^1–3^ for RNA-derived compounds, such as enzyme cofactors and ribonucleotide derivatives,^4,5^ have been discovered in recent years. These findings are consistent with the RNA World theory,^6,7^ which encompasses the concept that ancient metabolic processes and their regulation would have required many types of structured noncoding RNAs (ncRNAs) to perform various functions. As efforts to experimentally validate novel riboswitch classes have progressed, some observations revealed that many bacterial species also sense ligands whose biological relevance had previously been obscure. For example, the discovery of fluoride riboswitches^8,9^ has exposed the existence of numerous genes whose expression allows cells to overcome the stresses caused by toxic levels of this elemental anion.^9–11^ In other instances, RNAs corresponding to riboswitches for the bacterial second messengers c-di-AMP^12,13^ and c-AMP-GMP^14,15^ had been identified before their signaling molecules were known to exist.

The experimental validation of the function of guanidine riboswitches^16,17^ provides another prominent illustration wherein the investigation of a riboswitch candidate uncovers hidden biological processes. Three distinct structural classes of riboswitches have been found that respond to guanidine, called guanidine-I,^18^ guanidine-II,^19^ and guanidine-III.^20^ These RNAs control a diverse set of genes whose protein products act to protect cells from toxic levels of guanidinium ions. For example, a prominent set of genes associated with guanidine riboswitches codes for proteins implicated in guanidine degradation by first promoting guanidine carboxylation.^18,21^ Another commonly associated set of genes codes for proteins recently demonstrated to expel guanidine from cells.^18,22^ Strangely, it is not yet clear how cells become exposed to toxic levels of guanidine. Presumably they experience a metabolic state wherein guanidine accumulates as a byproduct of other metabolic processes.^16,18^

In our ongoing effort to find additional riboswitch classes and other structured ncRNAs, we identified nucleotide sequences corresponding to a widespread and well conserved sequence element called the *mepA* motif (**Figure 1A**). The name for this candidate ncRNA was chosen due to its frequent association with *mepA* genes, which commonly reside immediately downstream from the conserved features that define the motif. The *mepA* motif was initially uncovered by newly implementing a bioinformatics approach called ‘GC-IGR analysis’ method.^23,24^ This computational pipeline was used to comprehensively examine the genome of *Clostridium botulinum* Strain A Hall, which produced the initial representatives of the *mepA* motif. Ultimately, we detected 1275 representatives with non-redundant sequences distributed across six phyla: Actinobacteria, Firmicutes, Fusobacteria, Spirochaetes, Synergistetes and Tenericutes.

**Figure 1.**
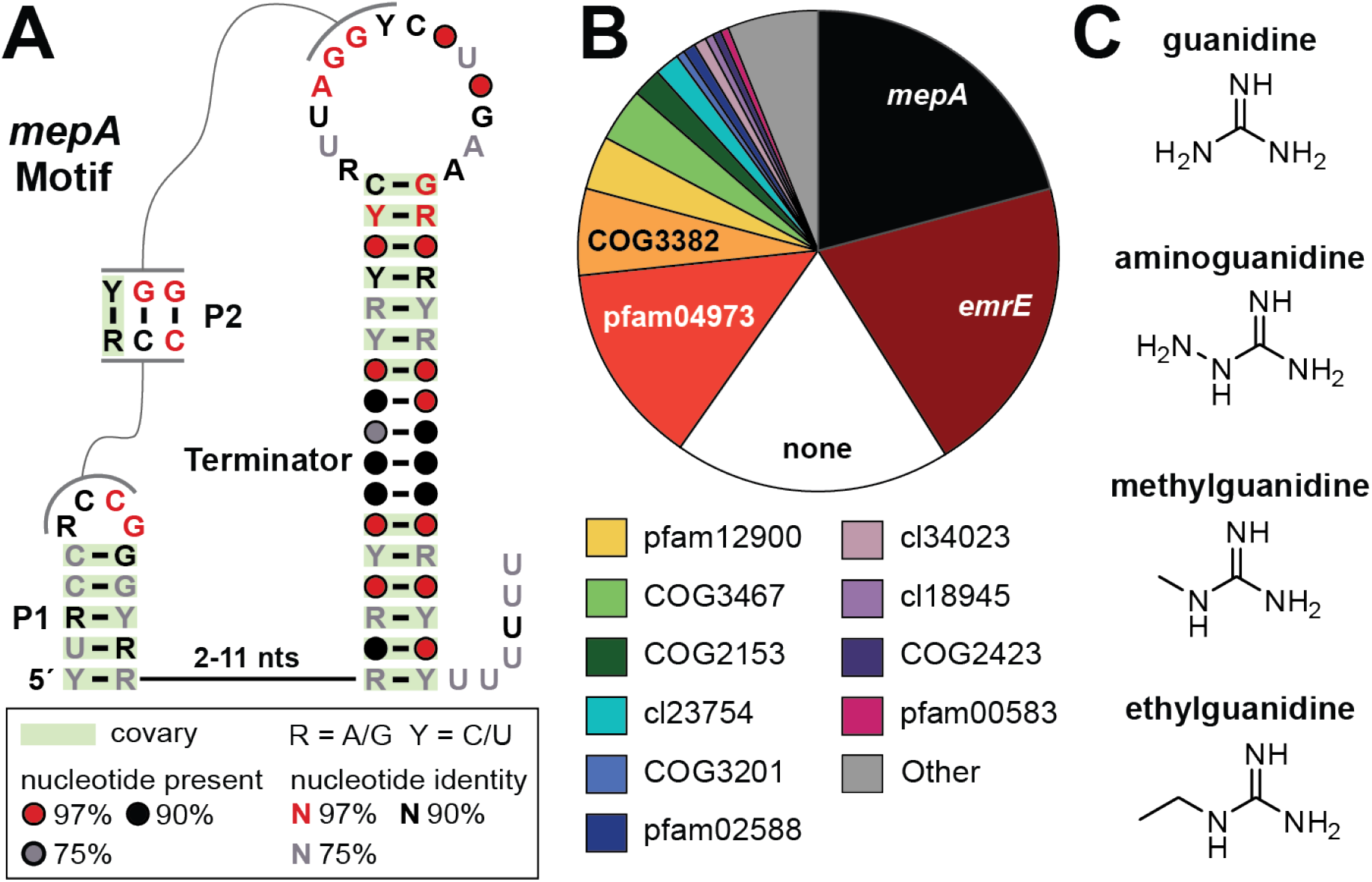
The *mepA* motif RNA associates with genes regulated by experimentally validated guanidine riboswitches. (A) Consensus sequence and secondary structure model of the *mepA* motif RNA based on 1275 homologous but non-identical sequences. P1 and P2 denote predicted base-paired stems, wherein P2 is incompatible with the formation of the predicted intrinsic terminator stem. (B) Genetic contexts of the *mepA* motif. Genes located immediately downstream of the motif are identified and reported (total of 1250). The difference between the number of adjacent genes recorded and the number of non-identical representatives is due to the fact that some representatives present in metagenomic sequence data lack sufficient read length to identify the neighboring gene. Additional details regarding the associated genes are presented in **Figure S1**. (C) Chemical structures of guanidinium and its analogs that exhibit the ability to be bound by a representative *mepA* RNA aptamer.

The diversity of sequences and broad phylogenetic distribution indicates that *mepA* motif RNAs serve an important biochemical role. The conserved features of this class (**Figure 1**) include two short base-paired substructures (P1 and P2) in addition to a typical intrinsic terminator stem.^25,26^ The most highly conserved nucleotide positions cluster both near the loop of P1 and the loop of the terminator stem, which would become proximal on the formation of the predicted P2 stem. A variable-length linker joins the right shoulder of P1 with the left shoulder of the terminator stem, which appears to be too short to permit the loops of these two hairpins as described above on the simultaneous formation of P1 and the full-size terminator stem. This architecture suggests that *mepA* motif RNAs might function as riboswitches wherein ligand binding to an aptamer domain modulates gene expression by controlling transcription via an expression platform employing a terminator stem. Specifically, we speculated that *mepA* motif RNAs are most likely to function as genetic “ON” switches wherein ligand binding stabilizes the formation of P2 to preclude formation of the full-length terminator stem.

Representatives of the *mepA* motif most commonly associate with genes (**Figure 1B**, **Figure S1**) that are known to be regulated by previously validated classes of guanidine riboswitches.^18–20^ As noted above, the most common gene association codes for MepA/MATE-like proteins, which are annotated as multidrug efflux proteins. Previous findings^18,22^ demonstrated that the biological function of certain efflux proteins associated with guanidine riboswitches is to eject guanidine from cells, and that variants of these proteins likely emerged later in evolution to function as antibiotics resistance factors.^22^ Additional genes that frequently occur near the *mepA* motif RNA include EmrE-like and B3/B4 domain proteins that are also commonly regulated by representatives of the known guanidine riboswitch classes.^18–20^ Therefore, we sought to evaluate the hypothesis that *mepA* motif RNAs function as members of a distinct riboswitch class for guanidine. In addition, we evaluated a series of guanidine analogs for ligand function, including those carrying modest chemical modifications (**Figure 1C**) that reduce affinity with representative aptamers from the known riboswitch classes for guanidine.

Two constructs based on the *mepA* representative from *C. botulinum* B1 strain Okra were prepared for analysis (**Figure 2A**). The 97 *mepA* construct carried 97 nucleotides from the natural sequence plus two additional guanosine nucleotides on the 5’ terminus to facilitate production by in vitro transcription. A shorter construct called 81 *mepA* was also prepared to weaken the terminator stem and increase the length of the single-stranded linker. This latter construct takes into consideration the hypothesis that the aptamer domain and the full-length terminator stem might form in a mutually exclusive manner, as would be expected if the RNA functions as a genetic “ON” riboswitch.

**Figure 2.**
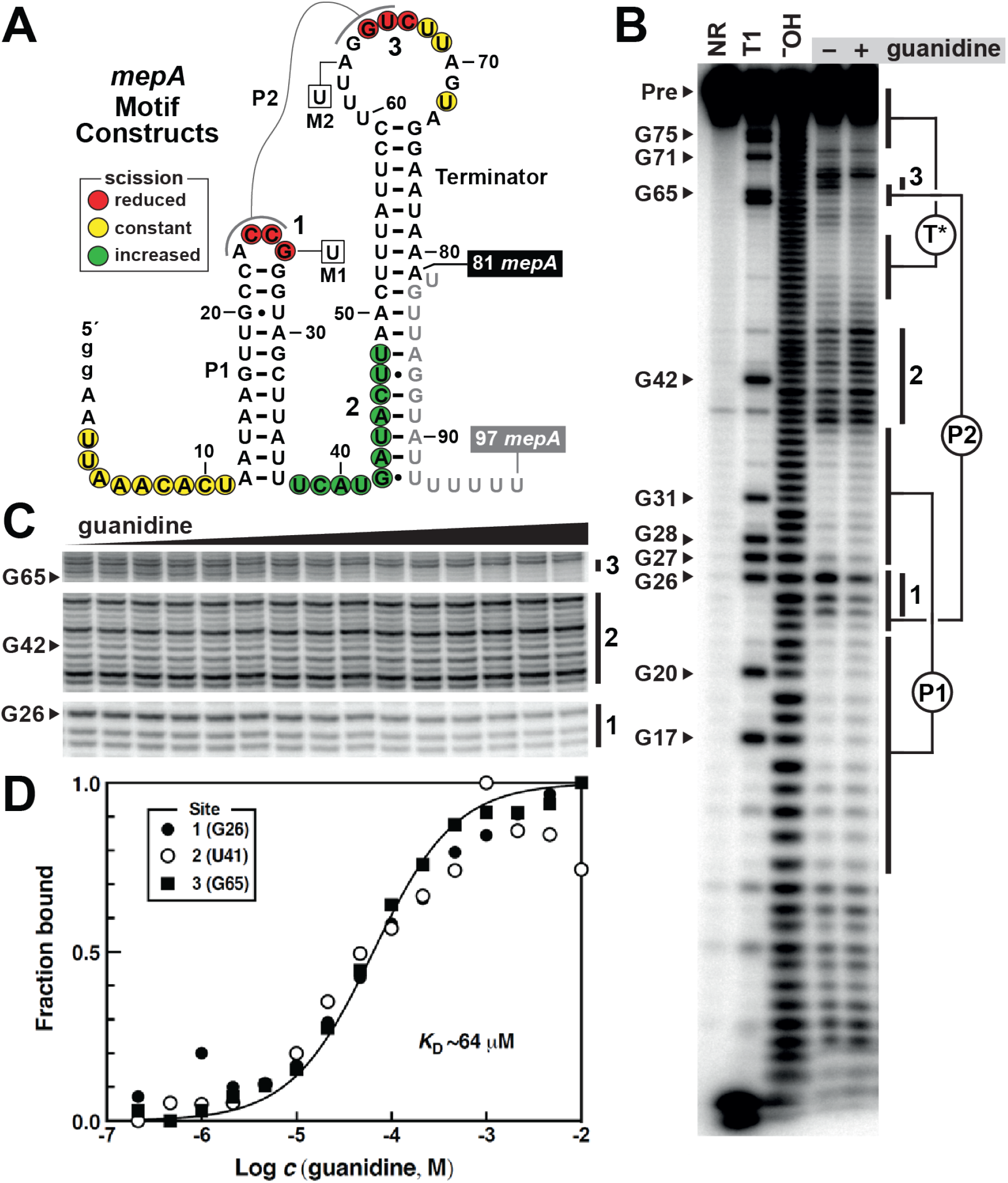
A *mepA* motif RNA construct undergoes a structural change when binding guanidine. A) Sequence and secondary structure model of a representative *mepA* motif RNA from *C. botulinum* B1 strain Okra. Two constructs examined in this study, 81 *mepA* and 97 *mepA*, have identical nucleotides on the 5’ side, but terminate at nucleotide positions 81 and 97, respectively. The two non-native guanosine nucleotides on the 5’ terminus (lowercase g letters) are not included in the numbering system used. Mutations present in constructs M1 (G26U) and M2 (A63U) are boxed. Results from in-line probing reactions using the 81 *mepA* RNA construct are derived from the data depicted in B. Colored circles indicate the sites undergoing substantial spontaneous strand scission in the absence and/or presence of guanidine. (B) Autoradiogram of the polyacrylamide gel electrophoresis (PAGE) separation of the products of in-line probing reactions. ^32^P-labeled 81 *mepA* RNA was subjected to in-line probing reactions in the absence (−) or presence (+) of 1 mM guanidine. Full-length 81 *mepA* precursor (Pre) RNAs present in the first three lanes were loaded without incubation (NR, no reaction), or were subjected to partial digestion with RNase T1 (T1, cleaves after G nucleotides) or alkali (^−^OH, cleaves after each nucleotide). Bands corresponding to strand scission after certain G nucleotides are labeled. In addition, regions corresponding to predicted P1, P2 and partial terminator (T*) base-paired structures are annotated. Regions exhibiting substantial modulation on the addition of guanidine are denoted sites 1, 2 and 3. (C) Portion of an autoradiogram of the PAGE separation of in-line probing reaction products generated near sites 1, 2 and 3 in the presence of various concentrations of guanidine ranging from 214 nm to 10 mM at every third log unit. (D) Plot of the fraction of 81 *mepA* RNA bound to ligand at various concentrations (c) of guanidine. Fraction bound values were derived from the data depicted in C, wherein the bands measured were G26 (site 1), U41 (site 2) and G65 (site 3). The line represents a theoretical curve for a 1:1 receptor-ligand interaction with a *K*_D_ of 64 μM.

Indeed, we failed to observe any evidence that the 97 *mepA* construct binds guanidine by using in-line probing assays^27,28^ (see below). In contrast, the 81 *mepA* construct exhibits robust structural modulation when 1 mM guanidine is added to binding assay reactions (**Figure 2B**). These assays take advantage of the fact that 3’,5’-phosphoester bonds in RNA can undergo spontaneous scission by the nucleophilic attack of a ribose 2’-oxygen atom on the adjacent phosphorus center. This reaction is dependent on the local structural context of an RNA molecule, wherein linkages in unstructured regions cleave more rapidly than those in highly structured regions.^27^ Thus, changes in the folded structure of an RNA on ligand binding cause changes in the pattern of RNA degradation products as observed by polyacrylamide gel electrophoresis (PAGE).

Two regions of 81 *mepA* observed to undergo suppression of spontaneous RNA strand scission in the presence of 1 mM guanidine (**Figure 2B**) are located in the loop of P1 (site 1) and the loop of the terminator stem (site 3) (**Figure 1A**). Nucleotides in or near sites 1 and 3 comprise the predicted P2 stem, suggesting that this *mepA* motif substructure is formed on ligand binding and therefore might be critical for aptamer function. In contrast, the nucleotides in the linker between the P1 and terminator stems (site 2) appear to be relatively unstructured in the absence of ligand, and these nucleotide linkages experience an increase in spontaneous scission when guanidine is present. This observation is consistent with our hypothesis that the ligand-binding aptamer conformation precludes formation of the full-length terminator stem. Finally, we note that nucleotide linkages preceding P1 and several linkages in the loop of the terminator stem that join poorly conserved nucleotides do not experience a change in spontaneous scission on the addition of guanidine, suggesting these nucleotides are not directly involved in forming the aptamer structure. Similar results were observed with an analogous construct based on the *mepA* motif representative from the bacterium *Lactobacillus brevis* (data not shown).

To establish the basic characteristics of the guanidine aptamer in the 81 *mepA* construct, we conducted in-line probing reactions using various concentrations of guanidine ranging from 214 nM to 10 mM. Quantitation of the intensities of certain bands from the resulting PAGE analysis of products (**Figure 2C**) allowed us to estimate the fraction of RNA bound to guanidine at each concentration tested (**Figure 2D**). The resulting plot indicates that the RNA binds guanidine in a 1-to-1 complex with an apparent dissociation constant (*K*_D_) of approximately 64 μM.

Similarly, binding assays were conducted with numerous guanidine analogs, which revealed that the 81 *mepA* RNA discriminates strongly against even close chemical derivatives. For example, at 1 and/or 10 mM concentration the methyl, amino, and ethyl derivatives of guanidine (**Figure 1C**) modulate the structure of the 81 *mepA* RNA in a manner essentially identical to that observed for guanidine as determined by in-line probing assays used to screen for ligand function (**Figure 3A**). However, additional assays using a range of ligand concentrations provided estimates of *K*_D_ values for these guanidine analogs (**Figure S2**), demonstrating that guanidine is bound by the aptamer 10-to 100-fold more tightly (**Figure 3B**).

**Figure 3.**
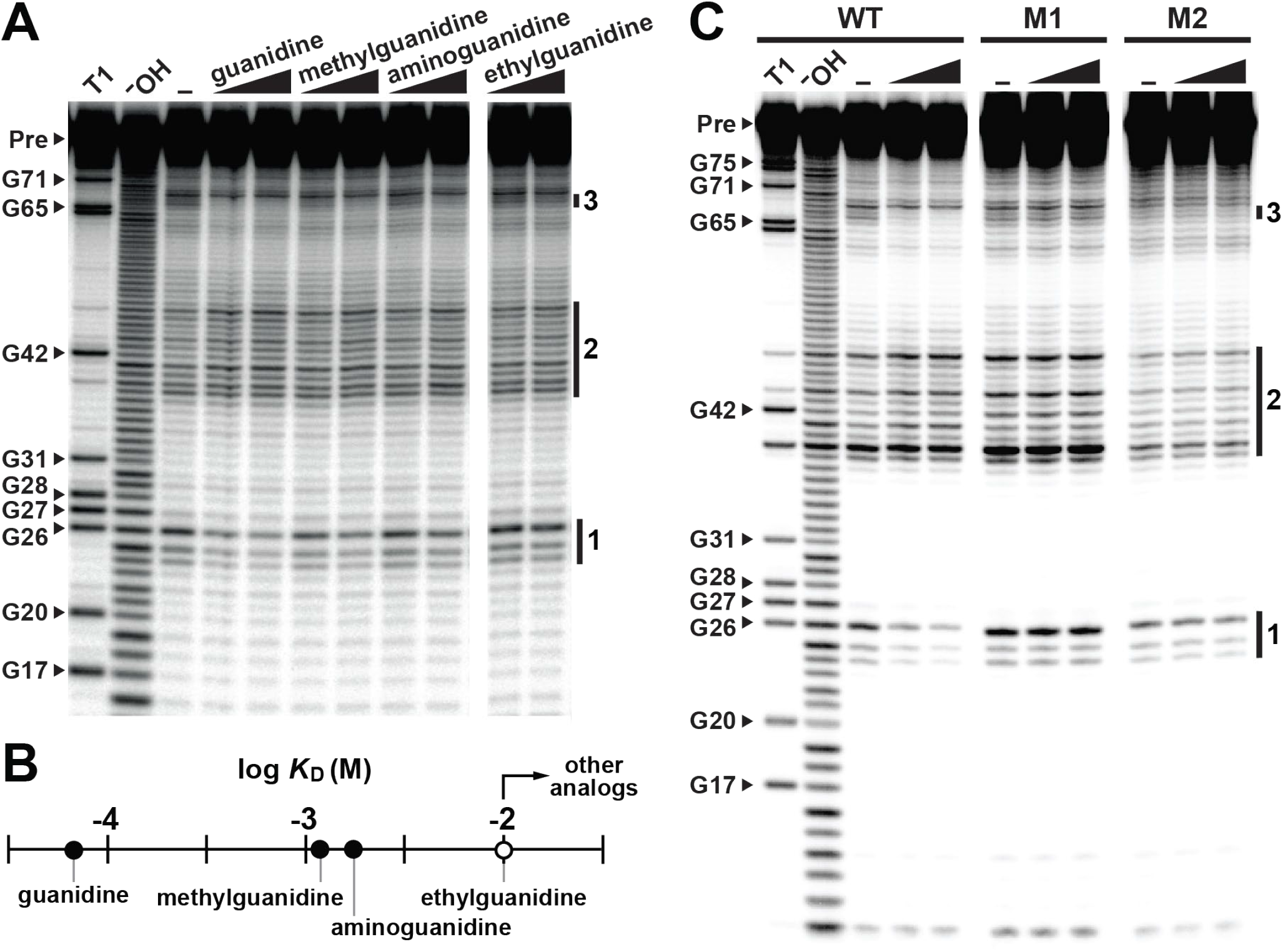
Requirements for ligand binding by the 81 *mepA* guanidine aptamer. (A) In-line probing analysis of ^32^P-labeled 81 *mepA* RNA in the presence of 1 or 10 mM of the ligands indicated. Other annotations are as described in the legend to Figure 2B. (B) Plot of the *K*_D_ values determined (filled circles) or estimated (open circle) for the ligands examined in A. See **Figure S2** for additional *K*_D_ analyses. Other guanidine analogs tested (1,1-dimethylguanidine; 1,3-diaminoguanidine; 1,1-dimethylbiguanidine; urea) exhibited no evidence of binding at 10 mM. (C) In-line probing analysis of radiolabeled WT (wild-type), M1 and M2 versions of the 81 *mepA* RNA construct. Guanidine concentrations tested were 0.1 and 1 mM. Other annotations, including the three sites of structural modulation exhibited by the WT RNA are as described in the legend to **Figure 2B**.

Additional analogs were also examined for evidence of binding (see the legend to **Figure 3B**), but all are rejected even when tested at concentrations of 10 mM (data not shown). The strong selectivity and affinity of the 81 *mepA* RNA construct for guanidine is very similar to the binding characteristics observed for members of the three previously studied guanidine riboswitch classes.^18–20^ The characteristics of these aptamers thus could be harnessed by cells to form precision riboswitches to regulate gene expression in response to elevated guanidine concentrations, despite the presence of numerous other metabolites naturally present in cells that carry a guanidine moiety.

Mutant versions of the 81 *mepA* RNA construct were also prepared to evaluate the importance of the highly conserved nucleotides in the loop of P1 or in the loop of the terminator hairpin. Specifically, mutant constructs M1 (G26U) and M2 (A63U) each carry a single nucleotide change (**Figure 2A**) at a position whose identity remains conserved in more than 97% of the known representatives (**Figure 1A**). As expected, these mutations independently disrupt guanidine binding, even when the ligand is present at 1 mM (**Figure 3C**). These results suggest that the conserved nucleotides observed by comparative sequence analysis are either directly forming the binding pocket for guanidine, or perhaps are required for *mepA* motif RNAs to adopt a structure that permits the pocket to fold properly.

The architecture of *mepA* motif RNAs is unusual compared to most known riboswitch classes. Most commonly, a contiguous aptamer domain is employed by a riboswitch, wherein this structure overlaps some portion of the expression platform to allow ligand binding to control production of either the full-length mRNA^4^ or the protein it encodes.^29^ However, *mepA* motif RNAs appear to carry a split aptamer, wherein the 3’ portion resides in the loop of the terminator stem. This creates a possible problem for aptamer formation because the two portions of the aptamer need to make contact to form the guanidine binding pocket. Again, this structural requirement is consistent with nucleotide covariation supporting the formation of a P2 stem (**Figure 1A**) and the guanidine-induced structural modulation evident from in-line probing assay data (**Figures 2, 3 and 4**).

**Figure 4.**
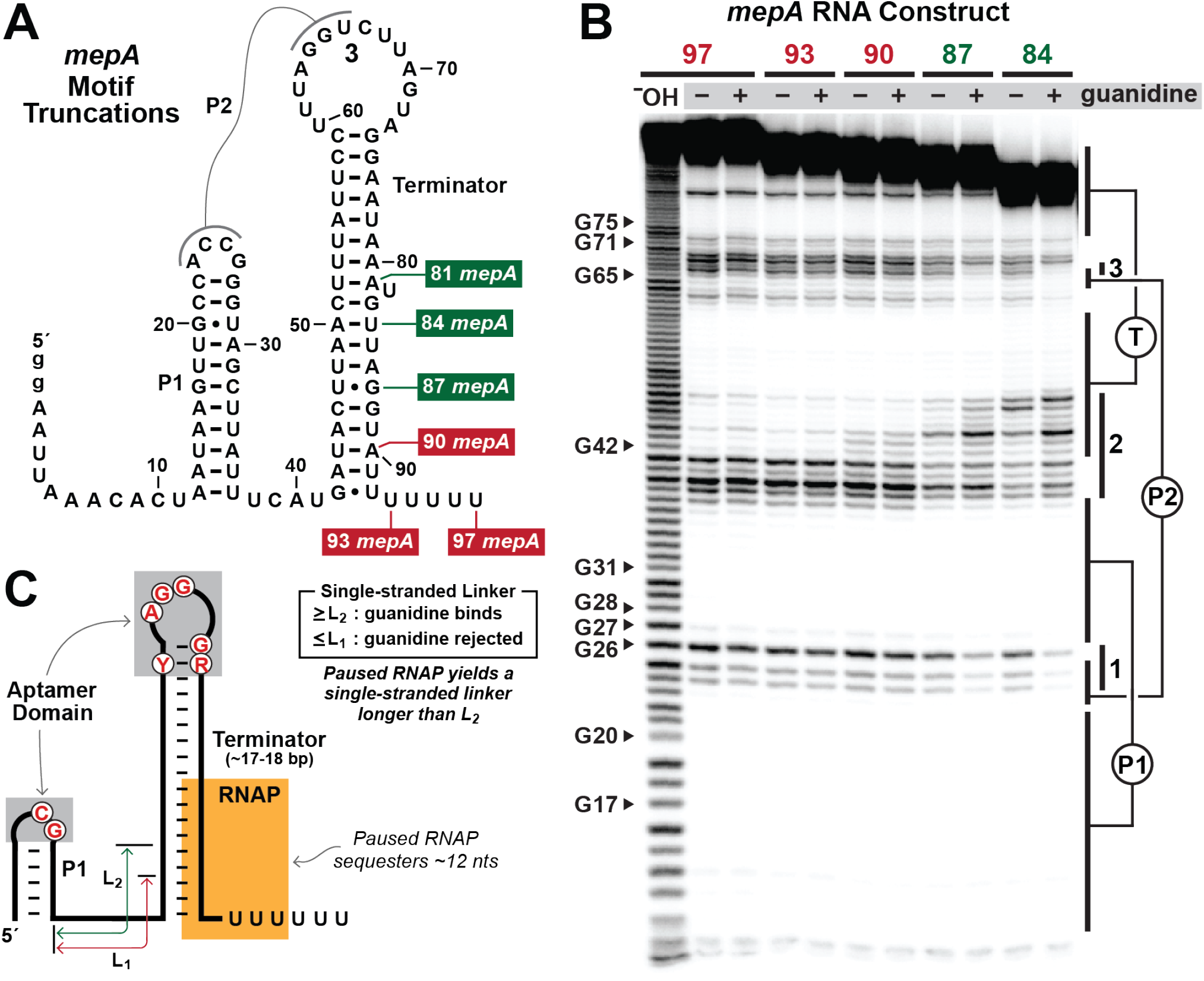
The mechanism of gene regulation employed by guanidine-IV riboswitches. (A) Truncated versions of the *mepA* RNA sequence from *C. botulinum* B1 strain Okra used to evaluate the importance of the terminator stem length on guanidine binding. (B) In-line probing analysis of various ^32^P-labeled *mepA* RNA constructs in the absence (−) or presence (+) of 1 mM guanidine. The bands corresponding to the terminator stem (T) are indicated. Other annotations are as described in the legend to **Figure 2B**. (C). Model depicting architectural features of *mepA* motif RNAs expected to be important for gene regulation by guanidine-IV riboswitches. The aptamer domain is likely formed by two regions of the RNA that carry highly conserved nucleotides (red letters). The length (L) of the single-stranded linker joining P1 and the terminator stem is proposed to be critical for the hairpin loops to make contact and form the guanidine aptamer.

This unusual architecture is predicted to create distance constraints that are coupled to each other in a manner dictated by the progression of RNA polymerase as it synthesizes the nascent mRNA. Specifically, for the parent *mepA* motif construct used in this study, the linker joining P1 and the left shoulder of the terminator stem could be entirely single-stranded until nucleotides corresponding to the right shoulder of the terminator stem begin to emerge from the exit channel of RNA polymerase. As the terminator stem begins to form base-pairs, the length of the single-stranded linker between P1 and the terminator becomes progressively shorter. Thus, we speculated that perhaps the length of the nascent terminator stem would dictate the ability of the two portions of the aptamer to make contact and form a guanidine binding site.

To examine this hypothesis, we prepared a series of 3-end deletion constructs (**Figure 4A**) based on the original 97 *mepA* RNA that encompasses the entire motif, including the run of U nucleotides that are characteristic of an intrinsic terminator device. As predicted, shorter constructs such as 84 *mepA* and 87 *mepA* exhibit guanidine-dependent structural modulation with patterns (**Figure 4B**) that are like those observed for the 81 *mepA* construct (**Figure 2**). However, longer constructs such as 90 *mepA*, 93 *mepA* and 97 *mepA* fail to respond to 1 mM guanidine. These findings indicate that as the length of the base-paired region of the terminator stem increases beyond a certain point, the split aptamer domains cannot merge to form the ligand binding pocket.

This data is consistent with a model for gene control (**Figure 4C**) wherein the ligand must bind to the nascent RNA before the terminator stem becomes too long to permit contact between the two parts of the aptamer. From data gathered using the *mepA* motif representative examined in this study (**Figure 4B**), we conclude that the single-stranded linker is too short (L1, **Figure 4C**) when the terminator includes nucleotide position 90. In contrast, the single-stranded linker is sufficiently long (L2) when the terminator stem included only up to nucleotide position 87. Intriguingly, bacterial RNA polymerases are known to sequester approximately 12 nucleotides of the nascent RNA transcript in the exit channel.^30–32^ Therefore, when RNAP is experiencing a transcriptional pause induced by the run of U nucleotides at the 3-end of the terminator structure, a portion of the right shoulder of the terminator stem remains sequestered in the exit channel.

Even if RNAP is paused near the end of this U-rich region, for example at nucleotide position 97, the nascent RNA transcript protruding from the protein complex only reaches approximately nucleotide position 86. Therefore, this paused state should still permit guanidine binding. Once guanidine binds, extending the base-pairing of the terminator stem should be disfavored because their complementary nucleotides in the single-stranded linker are structurally unavailable for interaction. This ligand-bound state thereby removes the thermodynamic incentive for base-pairs between the nascent RNA and its DNA template to be displaced in favor of the formation of additional base-pairs in the terminator stem, which is the widely accepted mechanism for this type of transcription termination.^25,26^

Similarly, we prepared a series of constructs based on the 81 *mepA* RNA with internal deletions within the putative single-stranded linker (**Figure 5A**) to further assess the effects of linker length on guanidine binding. In-line probing analyses reveal that the deletion of 6 nucleotides (construct Δ46-51) results in a functional aptamer, whereas the deletion of three more nucleotides (construct Δ43-51) prevents guanidine binding (**Figure 5B**). This additional data also sets constraints on the length of single-stranded nucleotides in the linker needed to permit ligand binding (**Figure 5C**). These findings combined with the length constraint noted earlier (**Figure 4**) indicate that the linker must have no less than 7 or 8 single-stranded nucleotides for the split aptamer portions to make contact and form a ligand-binding pocket.

**Figure 5.**
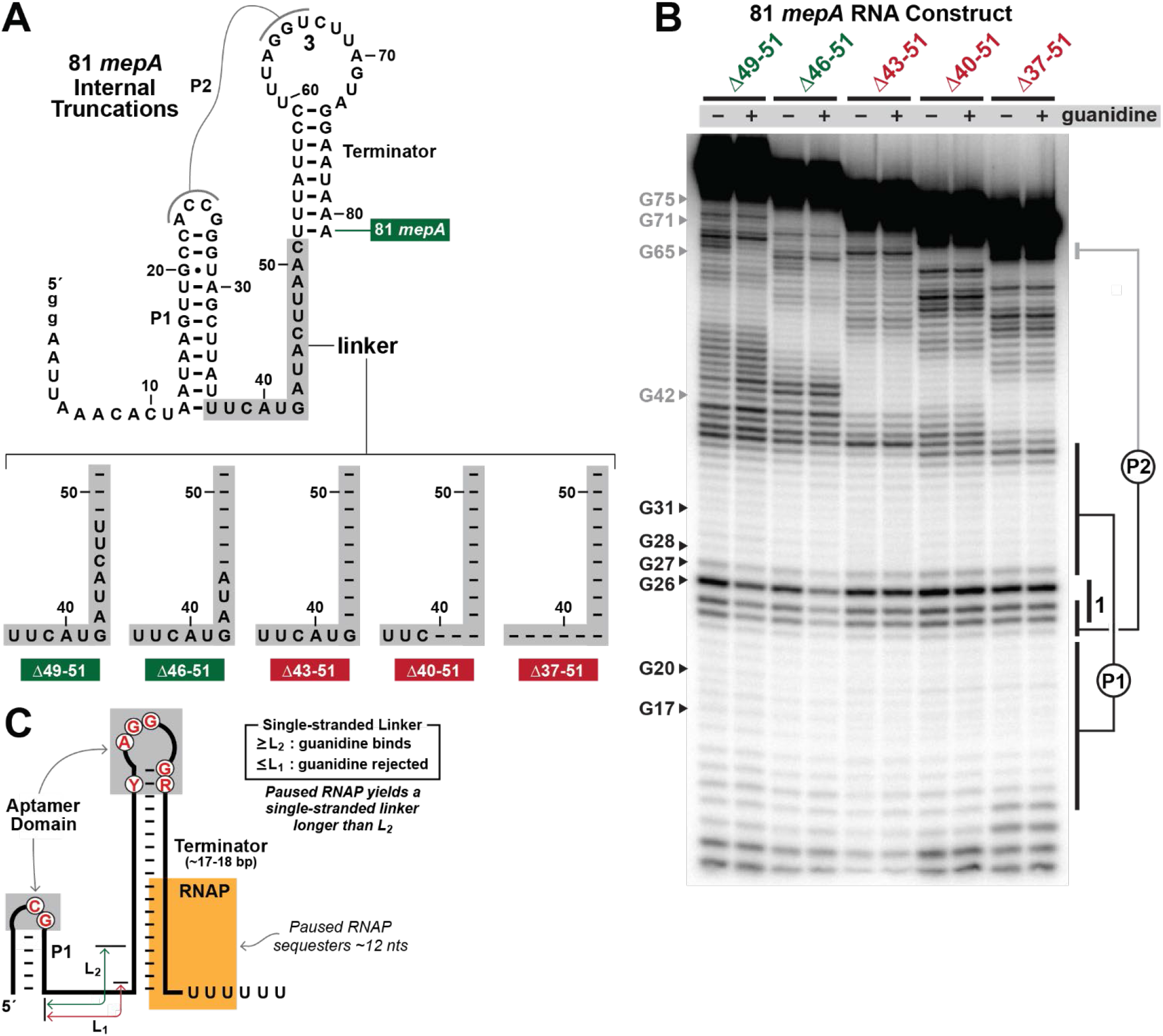
Internal deletions to the 81 *mepA* RNA construct support the proposed regulatory mechanism. (A) Internal deletions of the 81 *mepA* RNA were conducted in group of three nucleotides to create five additional constructs truncated within the proposed linker region (gray shading). (B) In-line probing analysis of ^32^P-labeled internal deletion constructs based on the 81 *mepA* RNA in the absence (−) or presence (+) of 1 mM guanidine. Annotations in gray apply only to the Δ49-51 construct. Other annotations are as described in the legend to Figure 2B. (C) Model depicting the key features of a guanidine-IV riboswitch, indicating the requirements for the length of the single-stranded linker. Other annotations are as described in the legend to Figure 4C.

Thus, given the bioinformatic, biochemical, and mechanistic characteristics of *mepA* motif RNAs, we conclude that they likely comprise as a fourth riboswitch class for guanidine. Thus, we propose naming this riboswitch class ‘guanidine-IV’. Our bioinformatics searches have uncovered 1275 non-redundant representatives of this newly found riboswitch class, which remarkably makes it more abundant in comparison to the numbers^1^ published for guanidine-I,^18^ guanidine-II,^19^ and guanidine-III^20^ riboswitches. Atomic-resolution structural models have been established for these previously reported riboswitch classes, demonstrating that RNAs can use diverse structures to selectively recognize guanidine.^33–37^ The biophysical analysis of guanidine-IV riboswitches promises to provide yet another glimpse at how RNAs can employ only four types of nucleotides to form precise binding pockets for this ligand. Taken together, this collection of riboswitch classes also provides further evidence that guanidine toxicity is widely experienced by many bacterial species.^16,17,38^ Interestingly, however, a biological source of the toxic levels of guanidine in so many species remains to be determined. Perhaps with the addition of the genes known to associate with guanidine-IV riboswitches, the expanded ‘super-regulon’ of guanidine-controlled pathways will provide some hints at the natural source(s) of this molecule. Regardless, advances are already being made on biological processes controlled by guanidine riboswitches,^21,39,40^ and guanidine aptamers appear to be promising RNA structures for use in synthetic biology.^41^

It is also interesting to consider why the guanidine-IV riboswitch class was not encountered in previous bioinformatics searches for novel ncRNA motifs.^8,12,42^ The consensus model exhibits extensive base-pairing and at least a modest number of highly-conserved nucleotides, which are features that typically designate strong ncRNA candidates during searches exploiting comparative sequence analysis algorithms. However, the consensus model is dominated by the terminator stem, which is usually ignored when ranking ncRNA candidates because of its prevalence as a known genetic feature. In other words, the presence of a terminator stem immediately upstream of a coding region is very common because this structure is typically the transcription stop site for the preceding transcriptional unit in many bacterial genomes.

When we first encountered representatives of the *mepA* motif, they appeared to be unusual to us because the loop sequences of the terminator stem were remarkably well-conserved, which is unnecessary for the structure to serve as a simple intrinsic transcription terminator.^25,26^ Indeed, the architecture of the *mepA* motif was reminiscent of a recently reported riboswitch class for the thiamin pyrophosphate biosynthetic precursor HMP-PP, which appears to exploit highly conserved nucleotides in a terminator stem to selectively bind its ligand.^24,43^ Likewise, we predict that nucleotides in the terminator loop are involved in forming the aptamer structure required to bind guanidine.

Given that riboswitches such as the HMP-PP class and the guanidine-IV class are more difficult to identify by previous comparative sequence analysis algorithms, there might be many other classes hidden in existing bacterial genomic databases that exploit the loop of terminator stems to form an aptamer structure. The unique structural constraints placed on riboswitches that embed all or a portion of their aptamer in the loop of an intrinsic transcriptional terminator hairpin are also likely to place functional constraints on the riboswitch. Thus, we predict that most RNAs with this architecture will function as genetic “ON” switches wherein the ligand binding structure and the terminator stem will be mutually exclusive structures.

## MATERIALS AND METHODS

### Chemicals

All chemicals were purchased from Sigma-Aldrich. Enzymes and oligonucleotide sources are described elsewhere.

### Bioinformatic analysis of *mepA* motif RNAs

The *mepA* motif was initially identified through the application of the GC-IGR genome analysis process^23,24^ to the genome *Clostridium botulinum* Strain A Hall as previously described.^23^ Briefly, the bioinformatic search was conducted with the Infernal 1.1 software,^44^ which was utilized to search genomic DNA sequences (RefSeq version 80) and microbial environmental sequence collections as described previously.^41^ Hypothetical gene annotations were validated manually by conducting protein sequence homology searches with the NCBI Basic Local Alignment Search Tool (BLAST).^45^ Covariation features were predicted using the comparative sequence analysis algorithm CMfinder.^46^ The RNA consensus model was generated with R2R software^47^ and manually adjusted to generate the final figure.

### RNA oligonucleotide preparation

RNA oligonucleotides were prepared as previously described^48^ using the appropriate synthetic DNA (Keck Oligo Synthesis Resource) templates for *in vitro* transcription (**Table S1**). Template DNAs were hybridized in equimolar amounts with their complementary strand to form a double-stranded DNA template containing a T7 RNA polymerase promoter sequence. The subsequent transcription reactions were incubated at 37°C for 4 hours. The resulting RNA transcripts were separated by using denaturing (8 M urea) 10% PAGE. RNAs were recovered from the gel, dephosphorylated using calf intestinal alkaline phosphatase, (New England Biolabs), and subsequently 5’^32^P-labeled using [γ-^32^P]-ATP and T4 polynucleotide kinase (New England Biolabs) according to the manufacturer’s protocols.

### In-line probing assays

In-line probing reactions were conducted generally as described previously.^27,28^ Briefly, ^32^P-labeled RNAs (trace amounts) were incubated in the absence or presence of ligand candidates as indicated at room temperature in the presence of 20 mM MgCl_2_, 100 mM KCl, and 50 mM Tris-HCl (pH 8.3 at ~23°C). The reaction products were separated by denaturing (8 M urea) 10% PAGE and were visualized by using a Typhoon phosphorimager (GE Healthcare). As described previously,^48^ band intensities were determined and used to estimate the fraction of RNAs bound to ligand. Values were plotted relative to the logarithm of the molar concentration of ligand, wherein half-maximal binding represents the *K*_D_.

## ACKNOWLEDGMENTS

We thank members of the Breaker laboratory for helpful discussions. Aparaajita Balaji was supported in part by the National Institutes of Health Chemical Biology Training Grant (T32 GM067543). This work was also supported by NIH grants (GM022778 and AI136794) to R.R.B. Research in the Breaker laboratory is also supported by the Howard Hughes Medical Institute.

## Supplementary Information

**Figure S1.**
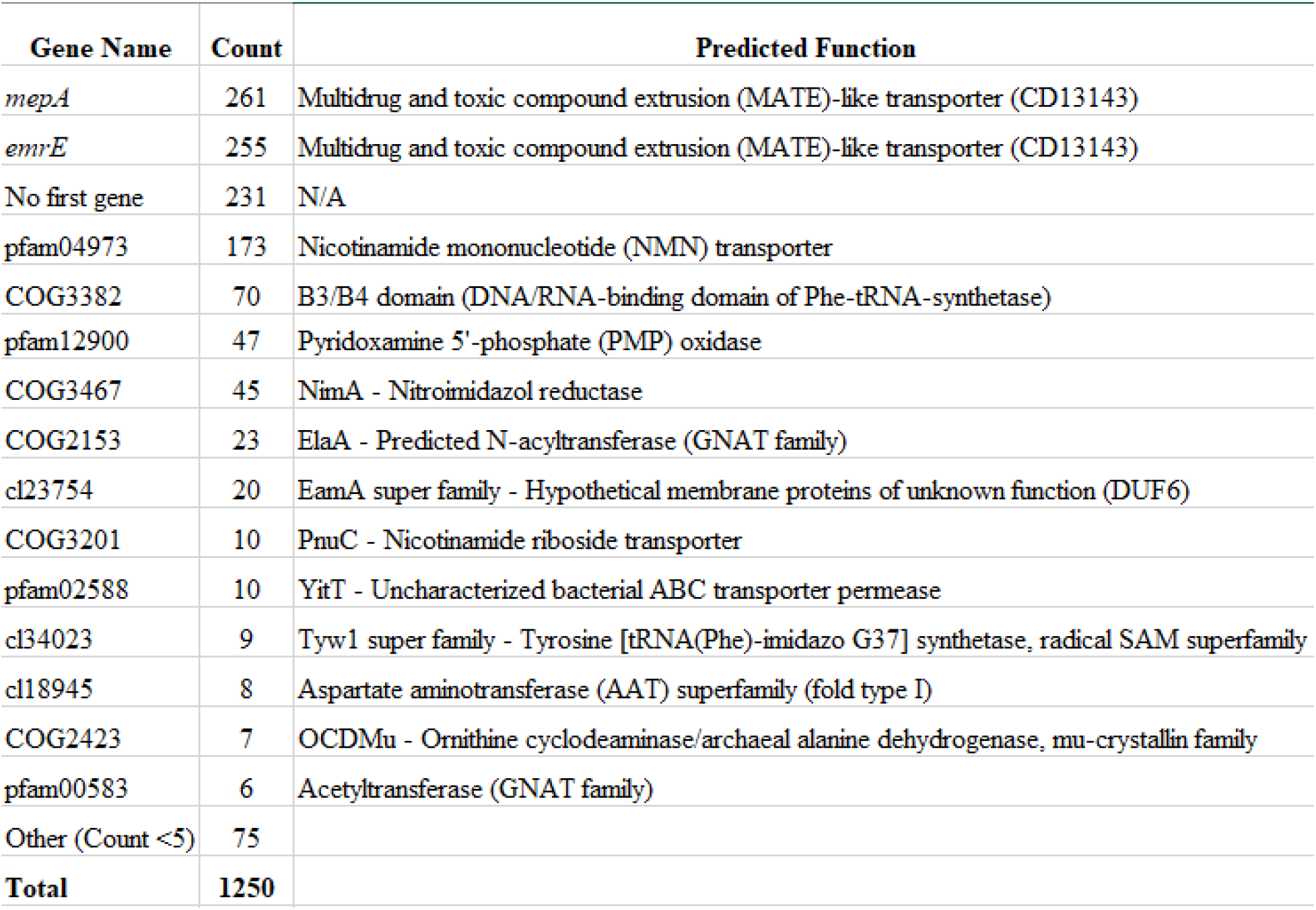
Existing annotations for genes associated with *mepA* motif representatives. N/A indicated not applicable.

**Figure S2.**
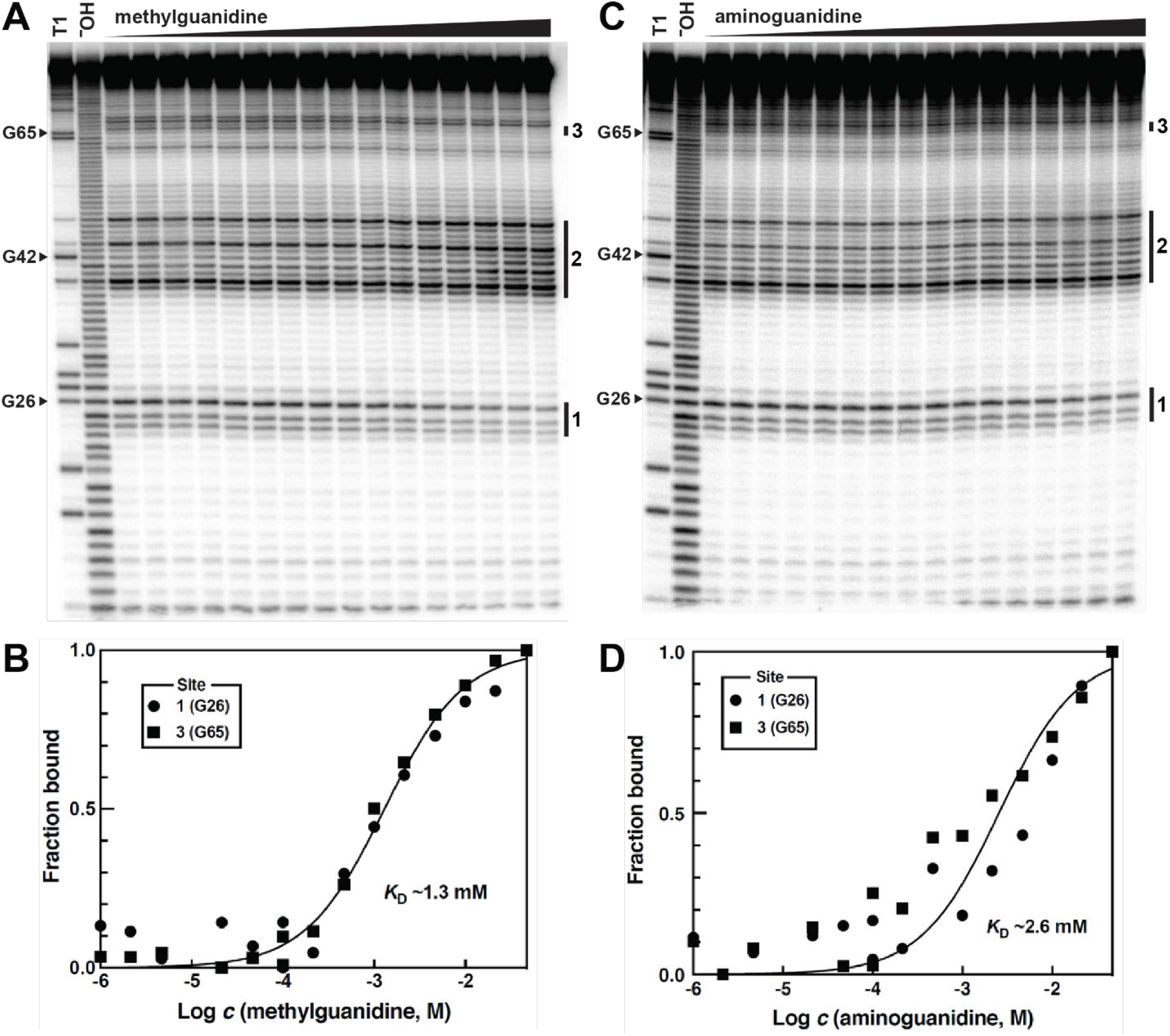
In line probing analyses to establish the apparent *K*_D_ values for guanidine analogs. N/A indicated not applicable. (A) The ^32^P-labeled 81 *mepA* RNA construct was subjected to in-line probing analysis with concentrations of methylguanidine ranging from 1 μM to 46 mM at each 1/3^rd^ log unit. (B) Plot of the fraction of 81 *mepA* RNA bound to ligand at various concentrations (c) of methylguanidine. (C) The analysis described in A using aminoguanidine. (D) KD analysis plot as described in B. Additional annotations are as described in the legend to **Figure 2**.

**Table S1.**
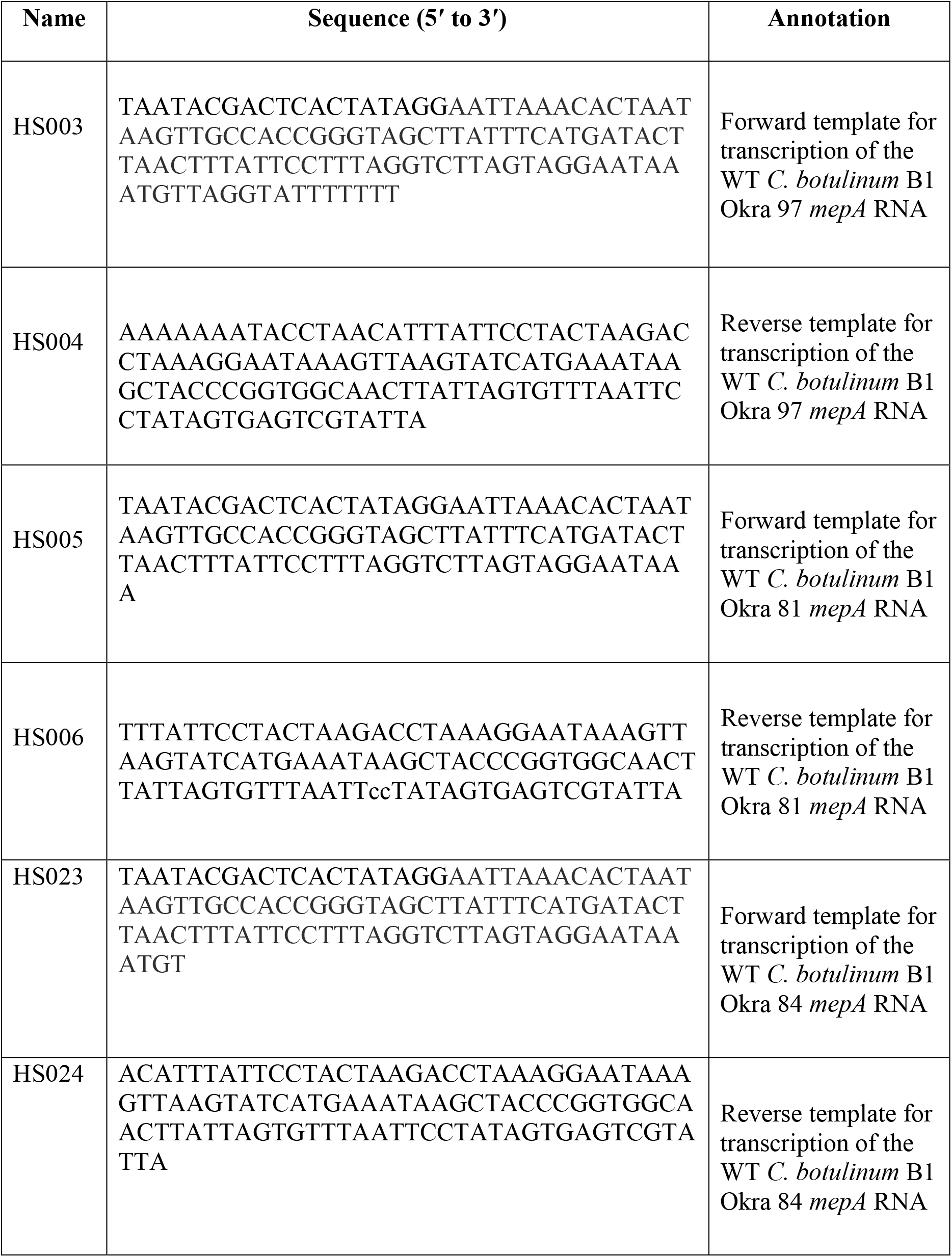
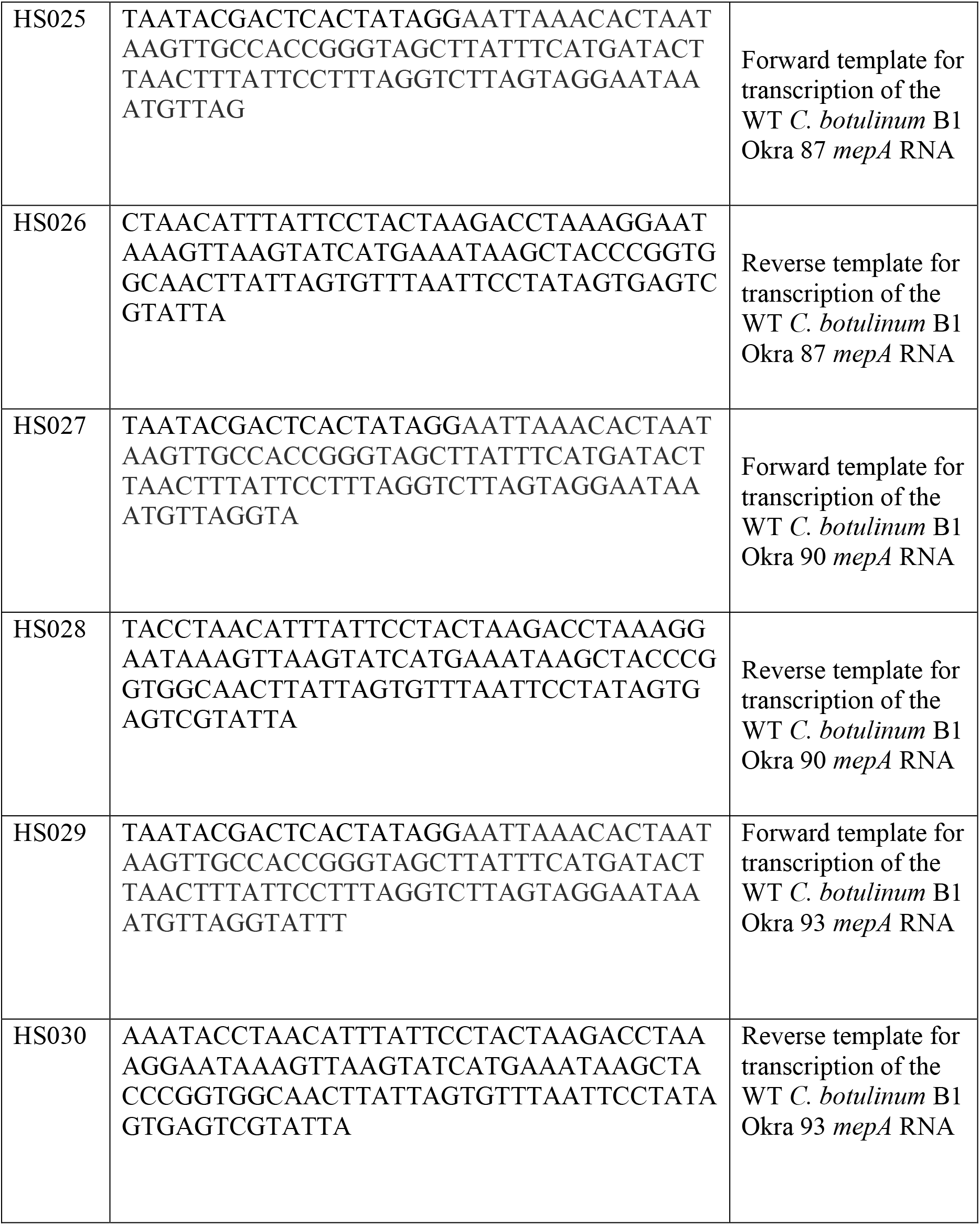
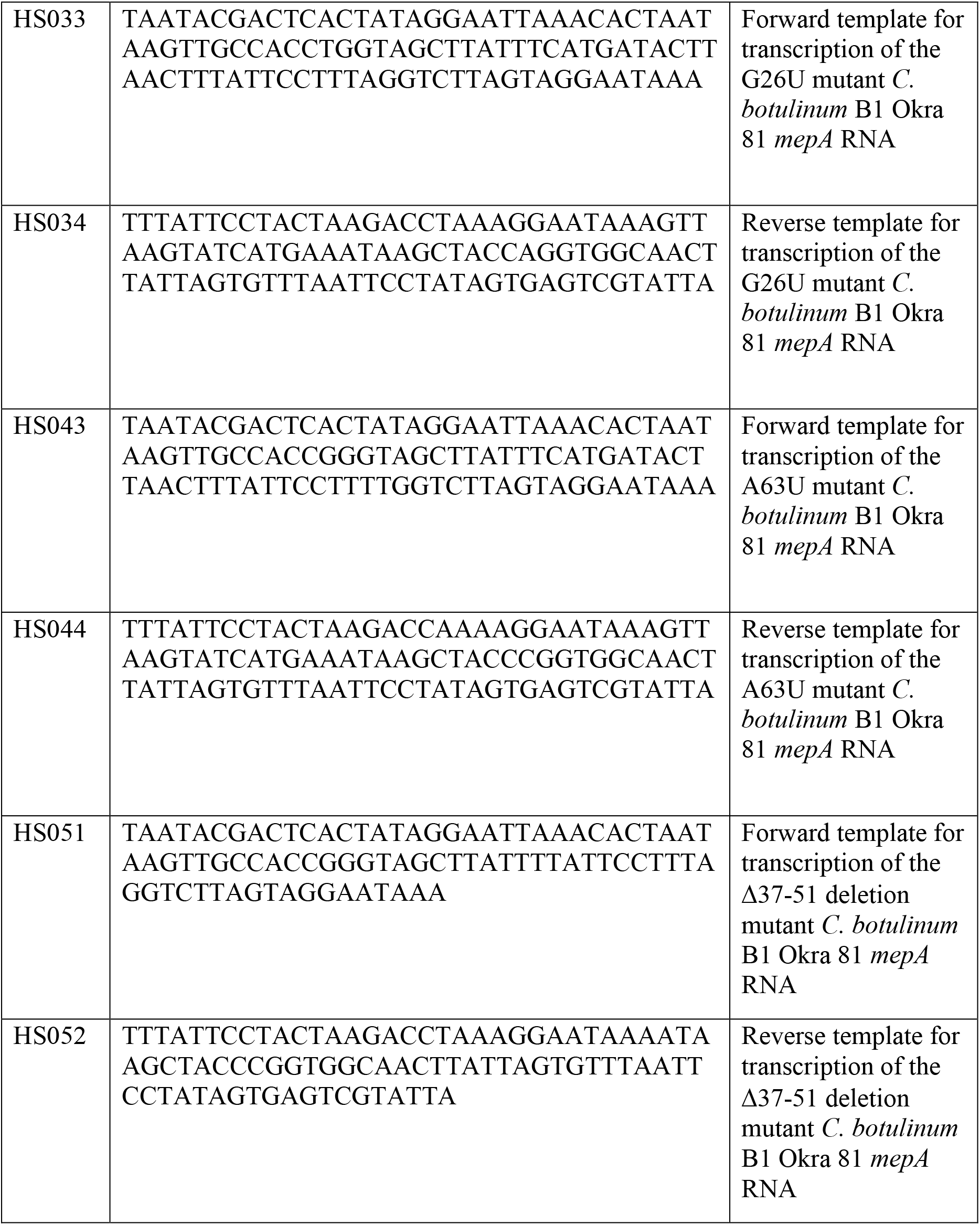
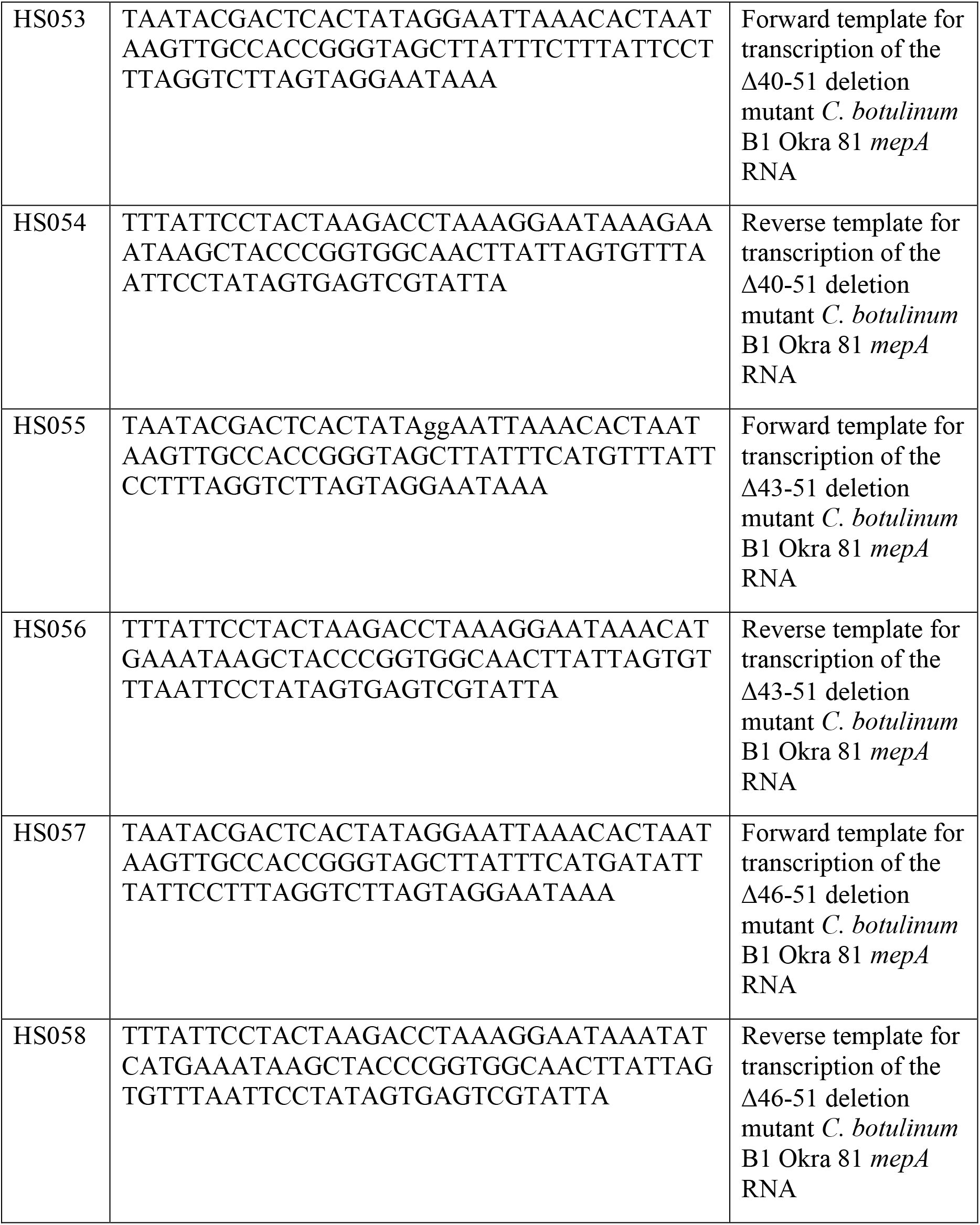
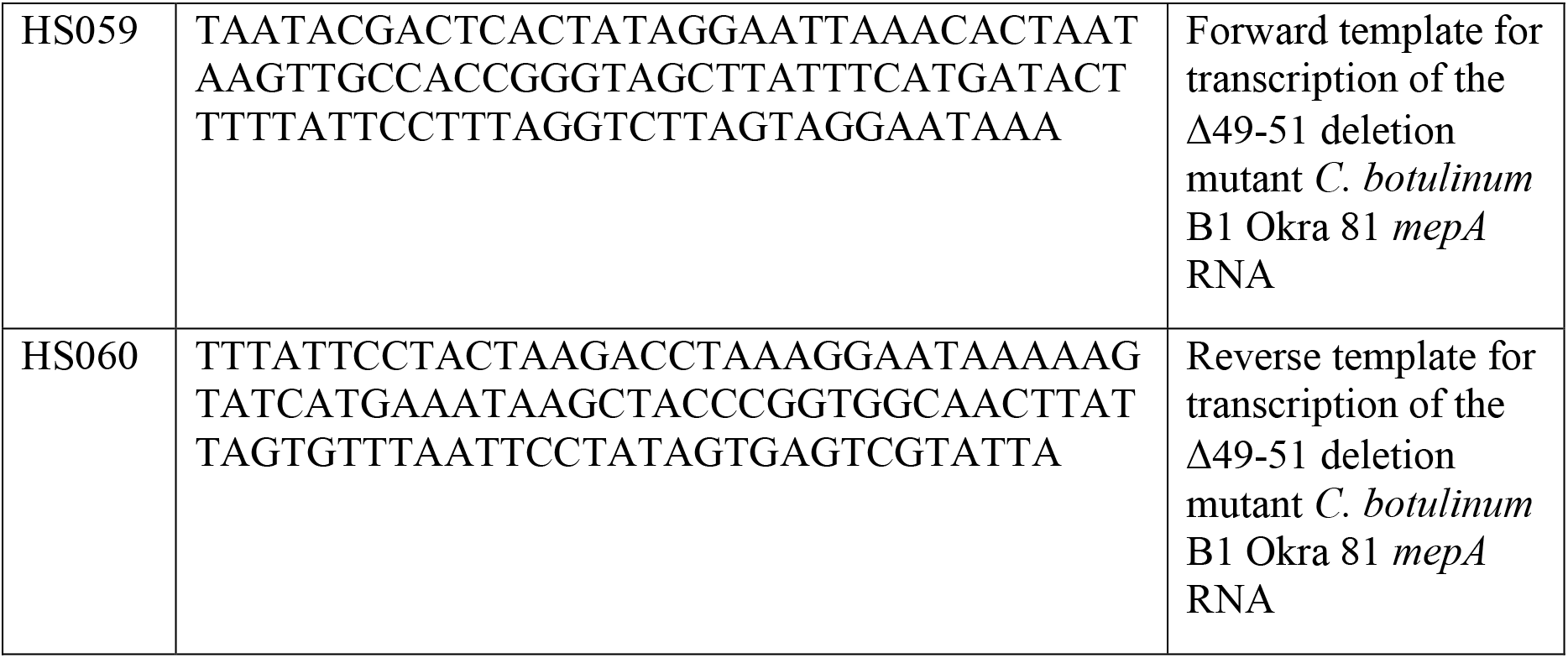
Sequences of synthetic DNA oligonucleotides used in this study.

